# Correlation of Remodeling Brain and Phenotype Behavior in the Comorbidity of the Chronic Obstructive Pulmonary Disease and Acute Cerebral Ischemia in Animal Model

**DOI:** 10.1101/469353

**Authors:** Natalia G. Plekhova, Boris I. Geltser, Sergey V. Zinoviev, Yulia V. Zayats

## Abstract

Cognitive impairment is one of the most common features in patients with COPD, and is associated with COPD severity and comorbidities. Development of relevant models of respiratory-cerebrovascular comorbidity in human diseases is an important task of experimental medicine. The purpose of the research consisted in determination of the morphological criteria brain condition and their correlation with behavioral phenotype of animals in the experimental comorbidity of the chronic obstructive pulmonary disease (COPD) and acute cerebral ischemia (ACI). Modeling of COPD on the basis of the combination of inducers, reproducing a proteoclastic degradation of lung tissue and systemic inflammation, and modeling of ACI by the suture middle cerebral artery occlusion with to mimic ischemia condition were used. Comparative histological study of the brain, neurological and behavioral phenotype of animals was conducted. It has been shown that in case of COPD and ACI comorbidity, formation of neurogliovascular microstructural complexes in brain is more pronounced than at animals with isolated form of disease, which was indicative of active adaptive transformation of neocortex. Significant disturbance of neurological and behavioral status of animals under the conditions of COPD and ACI comorbidity was correlated with the structural changes in the microvascular layer and neurons of brain. This study provides new insights about formation of neurogliovascular complexes with altered quantitative ratio in the vessels that was indicative of the presence of pericellular and perivascular edemas of the brain, and correlating of the these changes with the behavior of animals.

## Introduction

Chronic obstructive pulmonary disease (COPD) is a serious and highly prevalent condition associated with high mortality, and reduced quality of life. It’s disease of the lungs and airways, defined by expiratory airflow obstruction and, also, has several extrapulmonary manifestations and comorbidities, e.g. cardiovascular disease, osteoporosis, cachexia and skeletal muscle wasting (Hassel et al., 2014; Xu et al., 2016). Combination of COPD and ischemic stroke (IS) relates to the most frequent variants of “non-random” comorbidity – syntropy, determined by the presence of general pathophysiological patterns in development of these diseases (Geltser et al., 2018). The complexity of modeling COPD is associated with its multisystem manifestations, including cerebrovascular ones (Fricker et al., 2014; Perez-Rial et al., 2015). It was proved that in case of COPD, blood rheological properties deteriorate (Hawkins et al., 2015), atherogenesis progresses (Lankeit and Held, 2016), endothelial dysfunction develops, disturbance of carbohydrate and lipid metabolism occurs (Boukhenouna et al., 2018). Excessive proteolysis and oxidative stress against the background of chronic systemic inflammation produce effect on the molecular and cellular relationships in the nerve tissue, which results in the damage of neurons (Dodd, 2015). It is shown that dyscirculatory disorders in the cerebral circulation system in patients with COPD correlate with the intensity of cognitive dysfunction (Panagioti, 2018). It was found out that moderate hypoxemia is accompanied by an imbalance of perceptive and motor reactions and abstraction in 27 % of COPD patients, and in case of severe form of the disease this figure is as high as 62 % (Doehner et al., 2011). COPD is attributed to the most important factors of vascular comorbidity, which is manifested in development of «pulmonogenic» arterial hypertension, which is a leading cause for the development of the cerebrovascular pathology (Nobili et al., 2011; Wackera and Jorresb, 2016). It is noted that the incidence of IS in COPD patients is 20 % higher than among the general population (Haeusler et al., 2015). The increased risk of thrombotic events against the background of arterial hypoxemia and hypercapnia and hemodynamic instability produce a considerable effect on the IS occurrence rate (Morgan et al., 2017).

One of the top trends in fundamental science is experimental modeling of different variants of comorbidity in somatic pathologies (Traub et al., 2014; Bang et al., 2016; Huang et al., 2016). In these cases, creation of the relevant models of such states poses a certain problem. The study of COPD and IS comorbidity mechanisms, understanding the role and place of the respiratory and cerebrovascular syntropy in the pathogenesis of these diseases creates the prospects for implementation of the new strategies of individualized therapy.

The aim of the present study was the determination of the morphological criteria of the brain status, and their correlation with the behavioral phenotype of animals during experimental comorbidity of COPD and acute cerebral ischemia. We used a rat model with introduced a solution of papain in a total dose of 480 mg and reproduced systemic inflammation by introduction of bacterial lipopolysaccharide, which to lead to remodeling of lungs and were testifying to the moderate form of COPD (Geltser et al., 2018). After three months subject to established development of the disease the acute ischemic stroke was reproduced in animals. This allowed us to reproduce the comorbidity of these diseases and to study the changes in the brain and the behavior of animals.

## Results and Discussion

Modeling of COPD on the basis of the combination of inducers, reproducing a proteoclastic degradation of lung tissue and systemic inflammation, corresponds to the contemporary views on the pathogenesis of this disease (Fricker et al., 2014; Perez-Rial et al., 2015). The results of the microtomographic study of the respiratory system of the animals showed limitation of the inspiratory and expiratory air flow and the presence of hyperinflation. So, in rats with COPD, the volume of lungs on inspiration was 34% lower than in animals of control group (3224.6 ± 118.3 mm^3^ and 4318.6±193.9 mm^3^, respectively, p=0.032), and on expiration it significantly increased (2538.6 ± 79.3 mm^3^ and 2091.8 ± 88.7 mm^3^, respectively, p = 0.046). In addition, the increased airiness of the lungs was illustrated by a decrease in densitometric parameters calculated in terms of Hounsfield units: inspiratory (−674.2 ± 27.4 and −531.1 ± 21.1, respectively, p = 0.0293), expiratory (−468.2±24.3 and −377.8 ± 18.1, respectively, p = 0.0346). The development of respiratory failure and arterial hypoxemia was accompanied by the change of its functional indicators: reduction by 16 % of Sp0_2_ and increase by 42 % of respiratory frequency per minute. The first of them decreased by 16% (Sp0_2_ − 83.2 ± 2.53%; in the control 96.5 ± 2.1%, p = 0.047), the second - increased by 42% (respiratory frequency 118.6 ± 3.84 per minute, in the control − 83.7 ± 3.8, p = 0.0021). The visual assessment of the scans showed an emphysematous - pneumosclerotic version of the remodeling of the lung tissue associated with its elastopolytic destruction and chronic inflammation. The pronouncement of visual changes, as well as the quantitative evaluation the indicators for remodeling of lungs were testifying to the moderate and severe form of COPD.

Modeling of ACI in part of animals has lead to irreversible disorder of cerebral circulation and their death. In a group with combined pathology, the mortality was maximal and amounted to 80% on the 5-7th day, while in the group of animals with isolated ACI it amounted to 65 %. In the rest of the groups of animals, the survivability was equal to 100% (Table 1). The most pronounced changes of the neurological status took place in the survived animals with the combination of COPD and ACI. They included drastic limitation of mobility, retardation, disorientation, coordination disorders, inability to keep balance on horizontal bar and to perform purposeful movement along it. In all animals of this group, oculomotor innervation disorders, small-amplitude or medium-amplitude tremor of head, forelegs or tail, muscle weakness of limbs were registered. In addition, in most of this animals decreased reflexes of startle, blink, and fright when clapping. The sum of the points characterizing the severity of neurological deficit in animals of this group was maximum 48 hours after occlusion of the common carotid arteries and significantly exceeded this figure recorded 24 hours after the ACI simulation. At the same time, after 72 hours, no noticeable changes in the neurological status were observed. In combined of COPD and ACI, the neurological deficit corresponded to severe damage to the central nervous system, and in an isolated ACI, it was moderate. This was illustrated by significant differences in the sum of points on the NSS scale in rats of groups 4 and 5 at 24 to 72 hours after the simulation of the ACI. In animals with COPD neurological changes were slightly pronounced and did not increase during the entire observation period. At the same time, their neurological status was significantly different from intact animals and was manifested by restriction of the search behavior and reduction of the distance traveled along the crossbar.

**Table 1.** The survival and neurological status of animals.

| Animal groups | Indicators of survival rats, % |  |  | Indicators of scales NSS, points |  |  |
| --- | --- | --- | --- | --- | --- | --- |
|  | 24 hour | 48 hour | 72 hour | 24 hour | 48 hour | 72 hour |
| Intact animals groups, n=10 | 100 | 100 | 100 | 0,24±0,032 | 0,21±0,016 | 0,18±0,01 |
| Falsely operated animals, n=10 | 100 | 100 | 100 | 0,37±0,012 | 0,28±0,013 | 0,24±0,07 |
| COPD modeling, n=20 | 100 | 100 | 100 | 1,7±0,083* | 2,1 ±0,12* | 1,9±0,085* |
| ACI modeling, n=20 | 70 | 50 | 35 | 5,7±0,24 *# | 7,1 ±0,48*# | 6,8±0,29*# |
| COPD and ACI, n=20 | 50 | 40 | 20 | 7,8±0,27*#Д | 9,3±0,31 *#' | 8,8±0,37*#' |

Note: \* - the differences compared to 1 gr., # - compared to 3 gr., ' - between 4 and 5 gr., $p < 0.05$ .

The behavior of animals with isolated COPD was characterized by a decrease in locomotor function compared with those of intact. This was manifested by a 25 % reduction in the distance covered, with a relatively stable indicator of vertical activity (Fig. 1A,B). In animals of this group was no decrease in research activity and spatial memory, but manifestations of anxiety were recorded, which was illustrated by an increase in the number of short-term (“anxious”) grooming acts, a 24% reduction in the time of a long (“comfortable”) grooming (Fig. 1D) and the level of indicators in the eight-arm radial maze test (Fig. 2). The behavioral reactions of animals with ACI were manifested by a significant reduction in the distance traveled (2 times relative to the control) and the time of vertical activity, which indicated a sharp limitation of the locomotor function (Fig. 1A,B). In addition, in the rats of this group the level of spatial memory indicators recorded in the OSEM test decreased 1.8 and 1.6 times (Fig. 2C,D) and reduced by 54% the time of long grooming (Fig. 1). Behavioral phenotype in animals with a combined COPD and ACI demonstrated drastic reduction in the level of indicators of the horizontal (3.5 times) and vertical (2.7 times) activity comparing with the animals with isolated ACI, as well as minimal values of time for long-term grooming and sniffing of holes (Fig. 1). In rats of this group, by the results of the sniffing test (OF), the most marked disturbances of the vegetative support of the activity and anxiety-phobia imbalance with a marked limitation of spatial memory (Fig. 2). In the eight-arm radial maze test, the maximum deviations of the indicators from the control values were found out (in 3.3 and 2.6 times).

**Figure 1.**
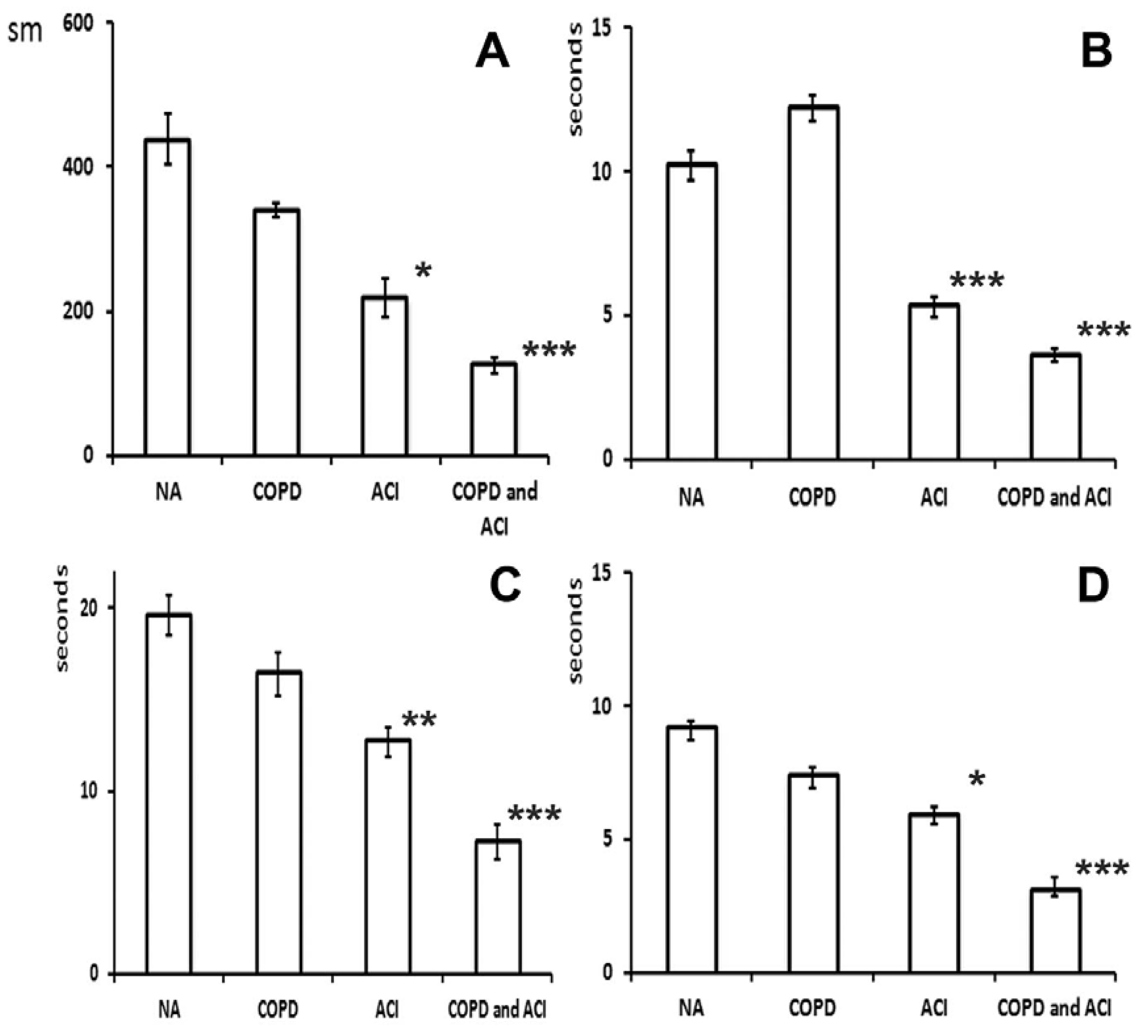
Indicators of Open Field (OF) test. A) distance traveled; B) total vertical activity; C) exploratory activity, time of sniffing holes; D) emotional state, number of long grooming acts. Data are mean±s.d.; *P<0.05; **P<0.01; ***P<0.001.

**Figure 2.**
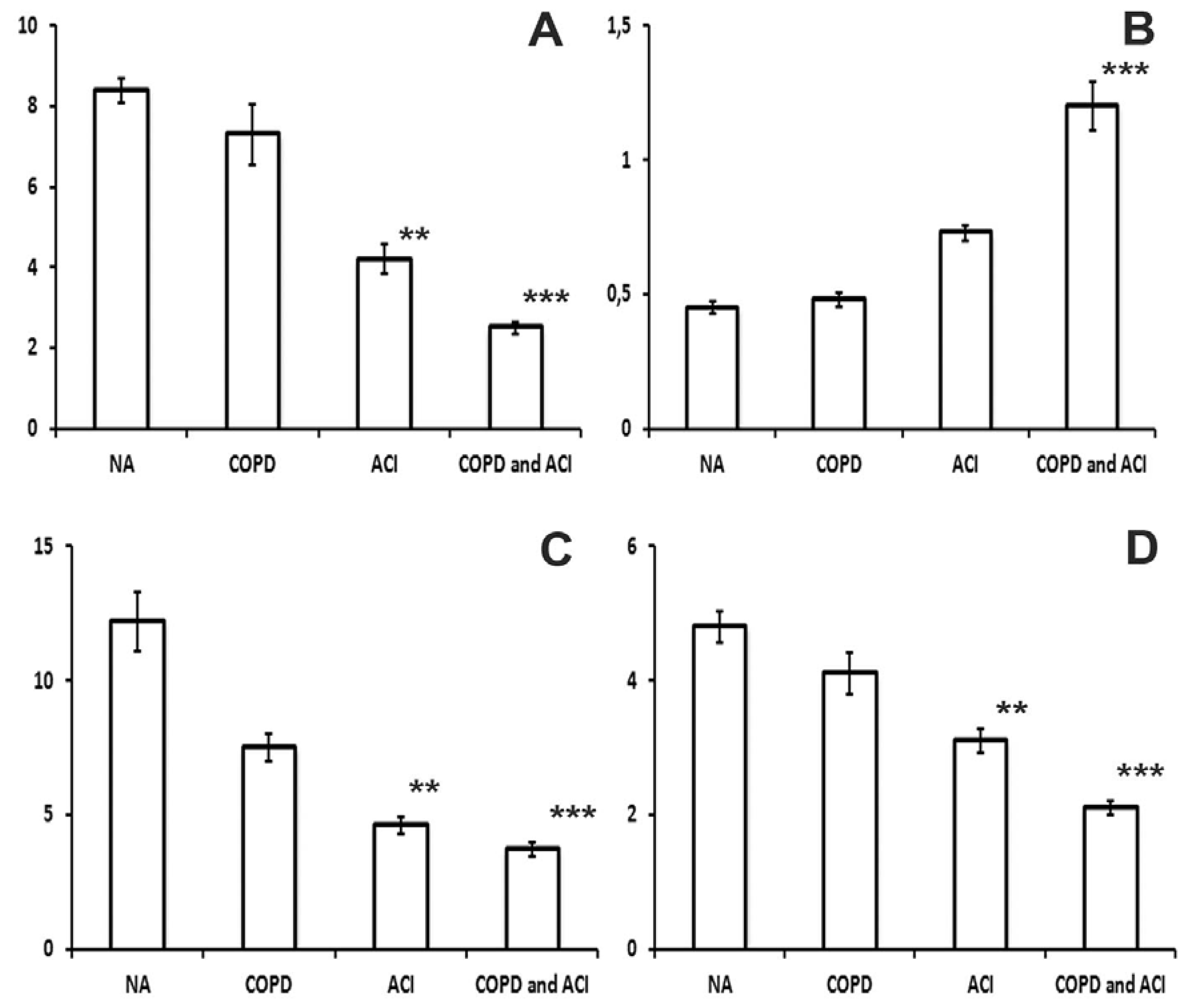
Indicators of Eight-arm Radial Maze (EARM) and O-shaped Elevated Maze (OSEM) tests. A) EARM, y-axis – the number of "correct" entrances to the sleeves, n; B) EARM, y-axis – the number of re-entrances to the sleeves, n; C) OSEM, y-axis – the number of exits to the non-protected sector, n; D) OSEM, y-axis – the time of the stay duration in the area, s. Calculation of the analyzed indicators was done within 5 minutes after adaptation of animals to the conditions of the experiment. Data are mean±s.d.; **P<0.05; ***P<0.001.

Despite the fact that it was possible to identify six layers represented by varying in size cells with predominance of pyramidal neurons (molecular, external granular, pyramidal, internal granular, ganglionic and polymorphous) in the cerebral cortex of rats in certain experimental groups, the general assessment of the nuclear cytoplasmic ratio allowed to identify various patterns of pathologic changes. In the neocortex of intact animals, in contrast to rats with induced pathology, normochromic neurons of symmetric shape with uniform chromatophilic cytoplasm, rounded nuclei and centrally positioned nucleoli were noted (Fig. 3A). In the brains of animals with experimental COPD, a considerable increase of the hyperchromatic neurons’ percentage was found out, which is a manifestation of hypoxia (Fig. 3B, Table 2). In rats with ACI, as compared to the animals with COPD more pronounced neuron changes were observed, which included emergence of a light space around the body due to the decrease of the cells size, which was indicative of the presence of pericellular edema (Fig. 3C). In addition to hyperchromatic neurons, the cells with vacuolated cytoplasm, hyperchromatic nuclei with unevenly distributed chromatin were noticed, which was indicative of its destruction due to necrosis. In the ischemia area, capillary hollowing was noted, while along the periphery of the necrosis locus, accumulation of cells was visualized (Fig. 3C). In animals with combination of COPD and ACI the total number of altered neurons was comparable with the brain of animals with isolated ACI, however, the predominance of the edematous cells was noted in all cytoarchitectonic layers of neocortex (Fig. 3D). In hyperchromatic nuclei of neurons, there were small, unclearly outlined nucleoli, adjacent to karyolemma. Morphometric analysis has shown the presence of intergroup differences between quantitative indicators of neuron and microglia dystrophy in animals with isolated ACI and in its combination with COPD, which was manifested by change of the ratio between the components of neuro-glio-vascular complexes (Table 3).

**Figure 3.**
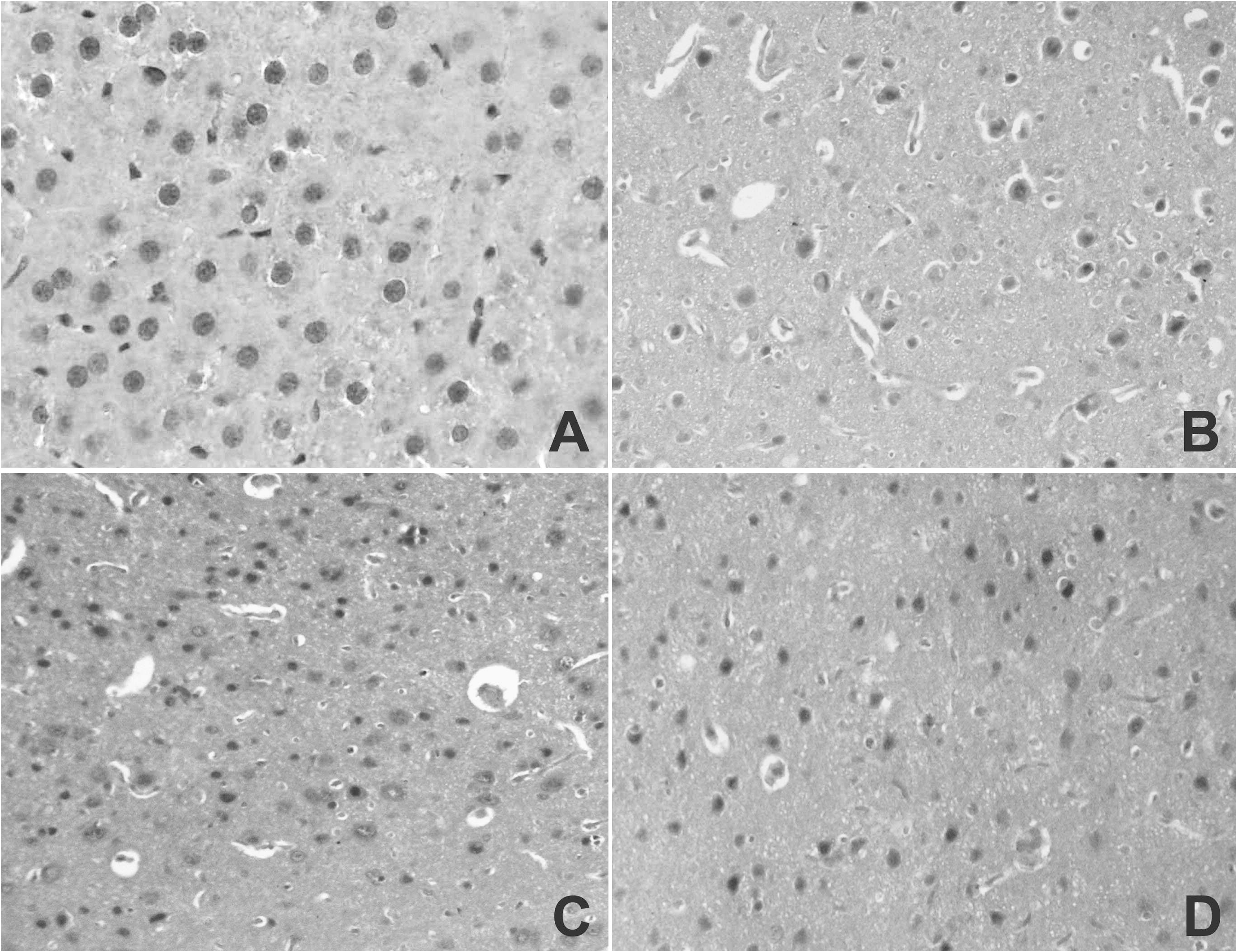
Neocortex of animals. A. Normochromic neurons with uniform chromatophilic cytoplasm in the cerebral cortex of the rats’ brains. Control. (400× magnification); Б. Hyperchromic neurons and luminal narrowing of the brain capillaries cerebral cortex in rats in the COPD. (200× magnification); В. Pericellular edema of neurons with hyperchromatic cytoplasm. Enlarged neurons with edematous cytoplasm with blurred boundaries, hypochromic nuclei and unevenly distributed chromatin and emptying of capillaries. Model of the ACI. (200× magnification); Г. The cells of edematous type by hyperchromic nuclei in the neocortex of animals with combination of COPD and ACI (200× magnification). Hematoxylin and eosin staining.

**Table 2.** The number of cells per 1 mm^2^ in Animals’ neocortex

| Group | Normochromic | Hyperchromic | Edematose |
| --- | --- | --- | --- |
| Control | 95,7±9,8 | 3,2±0,15 | 2,4±0,13 |
| COPD Modeling | 55,6±5,1** | 25,2±0,85** | 20,8±1,7** |
| ACI Modeling | 80,3±7,4 | 13,2±1,25** | 7,7±0,65 |
| Combination of COPD and ACI | 45,4±3,4** | 30,4±4,7** | 25,6±2,1** |

Remark: here and in Table 2, asterisks are used to denote the statistical significance
\* $P < 0.05$ , \*\* $P < 0.01$

**Table 3.**
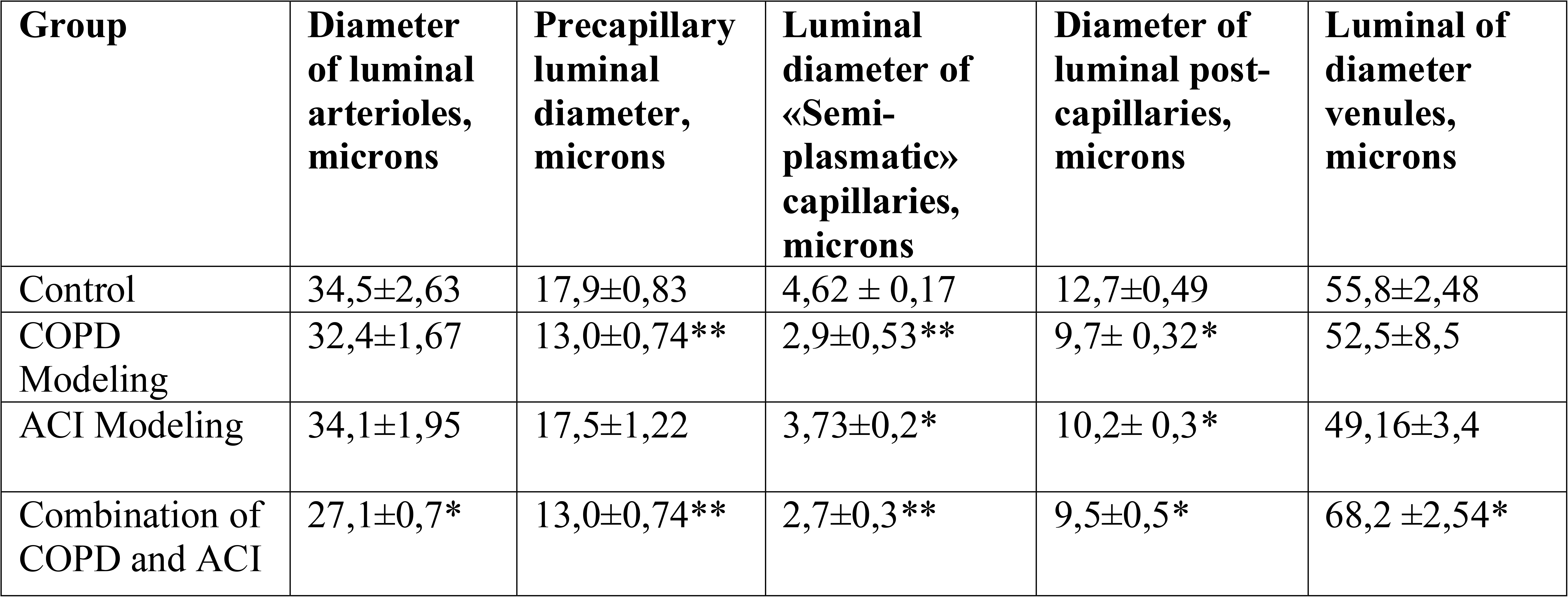
Morphometric indicators of the brain vessels animals

| Group | Diameter of luminal arterioles, microns | Precapillary luminal diameter, microns | Luminal diameter of «Semi-plasmatic» capillaries, microns | Diameter of luminal post-capillaries, microns | Luminal of diameter venules, microns |
| --- | --- | --- | --- | --- | --- |
| Control | 34,5±2,63 | 17,9±0,83 | 4,62 ± 0,17 | 12,7±0,49 | 55,8±2,48 |
| COPD Modeling | 32,4±1,67 | 13,0±0,74** | 2,9±0,53** | 9,7± 0,32* | 52,5±8,5 |
| ACI Modeling | 34,1±1,95 | 17,5±1,22 | 3,73±0,2* | 10,2± 0,3* | 49,16±3,4 |
| Combination of COPD and ACI | 27,1±0,7* | 13,0±0,74** | 2,7±0,3** | 9,5±0,5* | 68,2 ±2,54* |

During assessment of the vascular component of the brain, a significant spasm of precapillaries >5 mcm in diameter and capillaries <5mcm in diameter was noted in animals with isolated ACI and in the presence of COPD and ACI comorbidity (Table 3). In 95% of animals’ capillaries in this group, a one-row disposition of erythrocytes was noted with the presence of luminal narrowing in the areas filled with blood plasma (Fig. 4B,D). The shape of capillaries was altered, sinuous with incomplete filling and significant reduction of their lumen (Table 3). The walls of the capillaries were separated from the nerve tissue by a free space, by which the adventitia of the vessels was separated from the nerve tissue (Fig. 4A). Kernohan index of brain arterioles increased in animals with experimental comorbidity of COPD and ACI, whereas in case of isolated ACI no significant reduction of morphometric parameters of arterioles was observed.

**Figure 4.**
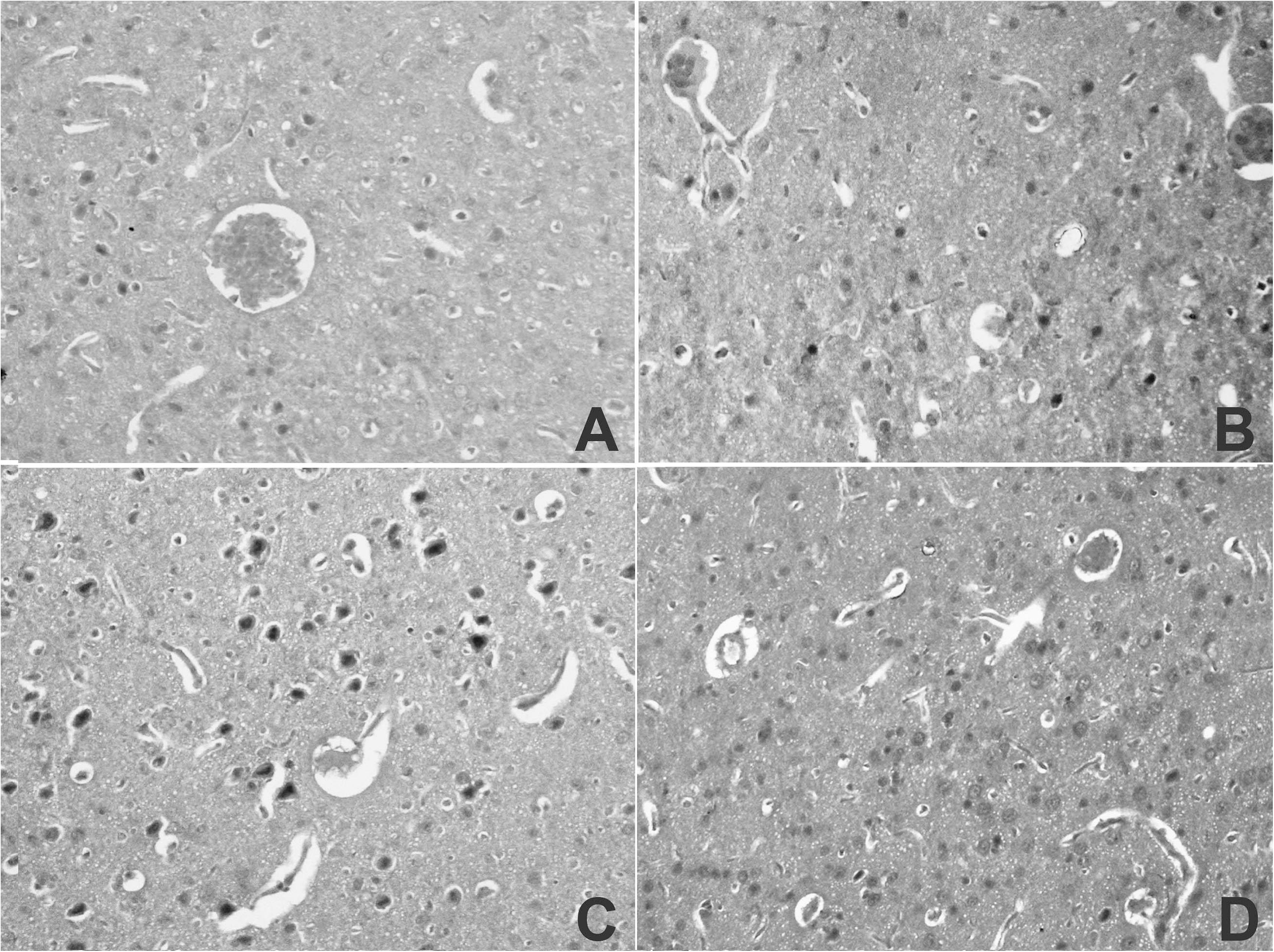
Vascular component of the animals brain in the modeling of the comorbidity of COPD and OCI. A. Wall of capillaries, delimited by free space from nervous tissue; B. Incomplete filling and reduction of the diameter of the lumen of meandering capillaries; B. Cells with vacuolated cytoplasm, hypochromic nuclei and unevenly distributed chromatin, along the periphery of the necrosis foci, cells accumulate; D. Single-rowed erythrocytes in the narrowed lumens of capillaries filled with blood plasma. 200× magnification. Hematoxylin and eosin staining.

So, the dependence of behavioral reactions animals on the type of pathologies has been established. With COPD on the background of the disease extrapulmonary manifestations, the behavioral disorders in animals were manifested by a moderate limitation of locomotor function, a slight increase in anxiety and preserved research activity and spatial memory. These data confirm the information about the correlation of the severity of cognitive disorders in patients with COPD with the degree of blood oxygenation (Doehner et al., 2011). Tissue hypoxia that accompanies such a state affects the structure of the brain and manifests itself at the cell level with a change in the level of neurotransmitter synthesis and neuroplastic damage.

Incomplete global cerebral ischemia, caused by the irreversible bilateral occlusion of the common carotid arteries, resulted in high mortality of rats and was accompanied by moderate neurological deficit in surviving animals. Already an hour after occlusion, there was a decrease in cerebral blood flow from 55 to 37 ml / 100 g / min and the development of acute cerebral ischemia, followed by a cascade of biochemical reactions inducing damage to neurons and neuroglia. Behavioral status was characterized by a significant limitation of locomotor function, exploratory activity, spatial memory, increased anxiety and the appearance of autonomic dysfunction signs. These changes demonstrated the known effects of ACI due to impaired intracellular metabolism of neurons as a result of a sharp restriction brain blood supply (Zhang et al., 2018).

The most significant changes in the neurological and behavioral status were occurred in the modeling of COPD and ACI in animals. There were a high mortality rate (80%) and maximal neurological disorders (ptosis, tremor, lethargy, disorientation). The behavioral reactions of these animals were characterized by minimal locomotor and exploratory activity, a sharp disturbance of spatial memory, high anxiety and increasing autonomic disbalance. Neurological and behavioral disorders in rats developed on the background of arterial hypoxemia and systemic inflammation associated with COPD, which created conditions for the initial cerebral discirculation and the more severe consequences of the simulated ACI. Neurological and behavioral disorders in rats developed on the background of arterial hypoxemia and systemic inflammation associated with COPD, which created conditions for the initial cerebral discirculation and the more severe consequences of the simulated ACI.

In the brain, partially autonomous neuro-glio-vascular microstructural complexes are being formed, the cellular components of which include neurons, glial cells (astrocytes), pericytes and endotheliocytes (Stepanov et al., 2017). In our study, such complexes were found predominantly in the brains of the animals with isolated COPD and ACI, which was indicative of the more marked adaptive transformation of nerve tissue in response to hypoxia. It has been shown that as the disease progresses, the dyscirculatory disorders in the cerebral pool increase due to the acceleration of the remodeling brain vascular system, increasing rigidity of vessels and further reducing blood flow velocity (Dodd, 2015; Richter et al., 2013). These changes are interrelated with the severity of ventilation disorders, arterial hypoxemia and hypercapnia (Richter et al., 2012; MacNee et al., 2013; Seeger et al., 2013). According to the voxel-based morphometry of the brain in the COPD decreases the volume of gray matter and most subcortical nuclei, which is in negative correlation with the partial oxygen tension in peripheral blood and in positive with the duration of this disease (Taki et al., 2013; Zhang et al., 2012, 2013; Wang et al., 2017). Behavioral phenotyping, as one of the methods of cognitive ethology, allows to evaluate the structure and dynamics of the animals’ reactions and characterizes their adaptation capability (Richter et al., 2014). Under the conditions of the experimental modeling of COPD and ACI, a pronounced disruption of the neurological and behavioral status of animals was correlated with the structural changes in microvascular bed (microvasculature) and brain neurons. Formation of neurogliovascular complexes with altered quantitative ratio in the vessels was indicatived of the presence of pericellular and perivascular edemas of the brain, which resulted in the altered behavior of animals (Dodd, 2012; Panagioti, 2014).

## CONCLUSION

The development of the relevant models of respiratory-cerebrovascular comorbidity is an important task of the experimental medical science. The presence of the pathophysiological patterns of the “non-random” combination of COPD and cerebrovascular disease is determined by the common risk factors and the systemic effects correlated with the chronic inflammation of the respiratory system. In the early stages of COPD, cerebral autoregulation is disturbed due to endothelial dysfunction in pre- and intracerebral arteries, which is manifested by a decrease of the dilatation reserve and increased of constrictor activity. Such changes are more pronounced in ACI. Still many questions remain about the causes and severity of the pathological process in the comorbidity of these diseases. The development of relevant models of such respiratory-cerebrovascular comorbidity allows objectively assessing the specific features of the neurological disorders and cognitive status in animals. Experimental modeling of such a state will solve many aspects characterizing the relationship between the morphofunctional state of the brain and individual components of cognitive disorders in comorbidity of COPD and ACI, and may also be useful for developing personalized programs for pharmacocorrection.

The many aspects characterizing the interrelationships between the morphofunctional status of the brain and certain components of cognitive disorders in the presence of COPD and ACI comorbidity remain unclear and they can be solved only by way of experimental modeling of such conditions.

## MATERIALS AND METHODS

### Animals

The experiment was conducted using sexually mature healthy male Wistar rats (4-6 months old; 200-250 g) were purchased from Puschino Laboratory Animal Research Center (Puschino, Russia) and housed in the animal room (25±3°C; 50% humidity; 12/12 h light and dark cycle) with free access to water and food. The animals were divided into 4 groups: 1 – reference group, intact animals groups (20 male); 2 – animals with COPD modeling, 3 – animals with modeling of acute cerebral ischemia (ACI, 20 male); 4 – animals with combination of COPD and ACI (20 male). All the potentially painful interventions in the conducted experiments, as well as euthanasia were carried out under a combined injection narcosis: Zoletil 0.003 mg/g («Virbac» France), Xilanit 0,008 mg/g (CJSC «NITA-PHARM», Russia, Saratov), solution of atropine sulfate 0,1% – 0.01 ml per 100 g. The experiments were performed according to the NIH Guide for the Care and Use of Laboratory Animals, the research design was approved by the local ethics committee of the Pacific State Medical University (№ 9, 18.10.2017).

### COPD animal model construction

The rat model of COPD was established as described in previous studies (Hassel et al., 2014; Machado et al., 2014; Brunnquell et al., 2014). Briefly, 20 rats in a 500 ml glass containment using an ultrasound device (UN-231, USA) were introduced a solution of purified papain (Upeen, Chine) of 10 mg/ml for 3 weeks in a total dose of 480 mg and reproduced systemic inflammation by intraperitoneal introduction of bacterial lipopolysaccharide (LPS; 200 mg/200 µl). The respiratory rate was controlled with the aid of physiological monitoring subsystem, saturation of arterial blood with oxygen (SpO_2_) was determined with the help of pulse oximeter Mouse Ox Plus (StarLife Science, USA). After 24, 48 and 72 hours of X-ray verification of COPD on the microtomograph «SKyscan-1176» (Bruker, USA) and post-operative recovery (model of ACI) the rats underwent a neurological test battery for three weeks comprising tests for exploratory activity, motor coordination and balance and spatial learning and memory.

### ACI animal model construction

To mimic ischemia condition the 20 rats were subjected to middle cerebral artery occlusion (MCAO) model for 2 h using the suture model, as has been described previously (Liu et al., 2017). Briefly, 20 rats were anesthetized with isoflurane (4% for induction, 1.75% for maintenance) during surgical procedures. Body temperature was maintained at 37.5 ± 0.5°C using a heating pad. The external carotid artery (ECA) and internal carotid artery (ICA) were exposed. A 4-0 silicone-coated monofilament nylon suture was inserted into the ICA via a cut on the ECA. Reperfusion was produced by gently withdrawing the suture out of the ECA. Successful surgery was further confirmed by tissue staining with 2,3,5-triphenyltetrazolium chloride (TTC). COPD and ACI animal model was established on the 20 rats after 72 hours of X-ray verification of COPD.

### Behavioral testing

Behavioral reactions were evaluated in the «Open Field» tests (OF), «eight-arm radial maze» (EARM), «O-shaped elevated maze» (OSEM) using «OpenScience» equipment (Russia). Video recording was done in real time with the use of digital videosystem EthoVision^®^ ХТ (USA). In the OF test, the locomotor function of animals (distance traveled and total vertical activity), exploratory activity (time of sniffing holes), emotional state (number of defecation acts, number of short- term and long grooming acts) were analyzed. In the EARM test, the parameters of spatial memory were estimated using the counting of correct and repeated inputs to the labyrinth. The level of anxiety of animals was investigated in the test of OSEM by the number of exits to the non-protected sector and the time spent in it. Calculation of the analyzed indicators was done within 5 minutes after adaptation of animals to the conditions of the experiment. The severity of neurological disorders was determined as the sum of scores according to NSS (Neurological Severity Scores), where light degree corresponded to the values from 1 to 3, moderate - from 4 to 6, severe – from 7 to 10.

*Brain histology* 10-14 days after the modeling of COPD and ACI the rats were anesthetized with isoflurane and killed. The samples were fixed in of 4% buffered formalin solution for >2 hours, they were paraffin embedded, cut and stained with hematoxylin, eosin and Nissl. The slides were examined under a light microscope with digital recorder (Olympus BX41/D25, Tokyo, Japan). An enumerated grid with each cell corresponding to a visual field under 200x magnification was applied and 10 fields chosen randomly and photographed for further investigation. Morphometric processing of images was carried out with the aid of the ImageJ 4.1 program.

*The statistical analysis* of the results was conducted with the aid of Statistica 6.0 (StatSoft, USA). The distribution normality was evaluated using Kolmogorov test according to the value of median (Ме) [25th; 75th percentiles], for evaluation of the credibility of the differences in comparison of two groups of variables, the The Mann-Whitney U-test and three non-prametric N-criterion Kruskal-Wallis tests (K-W) were used. The differences of indicators were considered to be statistically significant at p<0.05.

## Acknowledgements

A grant for the study was received from the Ministry of Health of the Russian Federation (government assignment).

## Author contributions

Conceptualization: N.G.P., B.I.G.; Methodology: N.G.P., S.V.Z., Yu.V.Z.; Validation: N.G.P., S.V.Z.; Formal analysis: N.G.P., B.I.G.; Investigation: S.V.Z., Yu.V.Z.; Resources: B.I.G.; Writing - original draft: N.G.P., B.I.G.; Writing - review & editing: N.G.P., B.I.G.; Visualization: N.G.P., S.V.Z.; Supervision: B.I.G.; Project administration: B.I.G.

